# Refget: standardised access to reference sequences

**DOI:** 10.1101/2021.03.11.434800

**Authors:** Andrew D Yates, Jeremy Adams, Somesh Chaturvedi, Robert M. Davies, Matthew Laird, Rasko Leinonen, Rishi Nag, Nathan C. Sheffield, Oliver Hofmann, Thomas Keane

## Abstract

Reference sequences are essential in creating a baseline of knowledge for many common bioinformatics methods, especially those using genomic sequencing. We have created refget, a Global Alliance for Genomics and Health API specification to access reference sequences and sub-sequences using an identifier derived from the sequence itself. We present four reference implementations across in-house and cloud infrastructure, a compliance suite and a web report used to ensure specification conformity across implementations. https://w3id.org/ga4gh/refget.

## INTRODUCTION

Since the release of the first human genome draft release (Lander *et al*., 2001), reference sequences have been central to genomic interpretation and to defining a baseline of knowledge upon which our understanding of biological systems, phenotypes and variation are based. The ability to interpret such data is essential for the delivery of genomics into the clinic (Birney *et al*., 2017). As precision medicine becomes mainstream in healthcare systems, organisations such as the Global Alliance for Genomics and Health (GA4GH) will help define interoperable standards, which will ensure the discovery and provenance of baseline knowledge.

Reference sequences currently suffer from two issues: sequence identity and non-standardised access. Genomic analysis methods, such as read alignment, typically use a FASTA-formatted collection of genomic sequences downloaded from a genome provider, *e*.*g*., an INSDC resource (Karsch-Mizrachi *et al*., 2018), Ensembl (Yates *et al*., 2020) or UCSC (Lee *et al*., 2020). These files use different names to refer to the same sequence. For example, chromosome 1 from the GRCh38 (GCA_000001405.15) release of Homo sapiens is also known as chr1 from hg38 (UCSC), 1 from GRCh38 (Ensembl) or CM000663.2 (INSDC). When it is critical to unambiguously identify an underlying reference sequence, it is better to use an identifier derived from the sequence itself, such as a cryptographic checksum digest. This method is employed in the CRAM format (Hsi-Yang Fritz *et al*., 2011), which uses a MD5 digest of a reference sequence ensuring the correct reference is used during read reconstitution. The European Nucleotide Archive (ENA) (Harrison *et al*., 2019) developed the CRAM reference registry (CRR) to retrieve reference sequences by an MD5 checksum digest derived from a reference sequence.

This manuscript describes a new application programming interface (API), called refget, which enables retrieval of full-length sequences or sub-sequences via an identifier, returns metadata associated with an identifier and maintains compatibility with the CRR. Our API operates over HTTP(s) and so is accessible in all main-stream programming languages. We also present four implementations of the refget specification deployed across in-house and cloud infra-structures and a toolkit to assess implementation compliance.

## RESULTS

### THE REFGET PROTOCOL

The refget protocol operates with a client providing a supported digest identifier with an optional linear or circular genomic coordinate range, specified as URL parameters or a Range header, via a HTTP(s) GET request. An implementation responds with a non-breaking stream of sequence characters. Users may request a metadata JSON document, which provides information about the sequence length, topology, known digests and any other known aliases. Finally, a client can request a JSON document of server capabilities allowing for clients to adapt to possible limitations of a server implementation. Implementations are not restricted to a single type of reference sequence to serve and therefore can provide DNA, mRNA, cDNA, CDS or peptide sequences. Should an implementation wish to provide a CRAM reference registry (CRR) compatible deployment they must mirror the reference sequences as found in ENA.

Refget defines three supported identifier algorithms: MD5, TRUNC512 and GA4GH Identifier. All three algorithms normalise sequences by stripping all whitespace characters and restricting to characters in the range A-Z. We chose this as a compromise between the methods and requirements employed by CRAM, ENA and the Variation Representation Specification (VRS).

MD5 is the default checksum algorithm used by the CRAM format’s M5 tag and hence the CRR and is supported to maintain compatibility with CRAM files. However, there are limitations to md5 and hash collisions are a known weakness. The occurrence of a checksum collision between non-identical sequences would be catastrophic to downstream reconstruction of sequencing reads. To mitigate this concern, we have used the sha512t24u identifier scheme (also known as the GA4GH identifier) as employed by the Genomic Knowledge Standards’ VRS standard (Hart and Prlić, 2020). In addition, we created a parallel format called TRUNC512, which represents the truncated SHA-512 digest as a hex string to maintain a similar representation to MD5. These schemes are described in Figure 1. However, sha512t24u is now the preferred representation due to its use in VRS. We applied TUNC512 (and therefore sha512t24u) to the MGnify (Mitchell *et al*., 2018) May 2019 protein database of ∼1 billion entries and found no collisions (see supplementary material).

**Figure 1.**
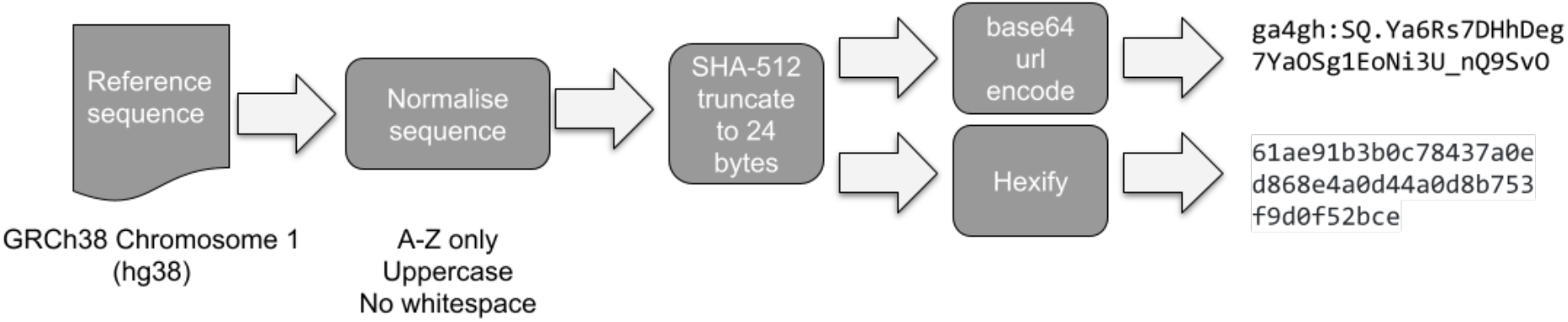
Summary of the sequence normalisation and algorithm used to generate checksum identifiers for TRUNC512 and GA4GH Identifier. All methods move through the same normalisation process but differ in their choice of checksum algorithm (MD5 vs. SHA-512)

### IMPLEMENTATIONS AND COMPLIANCE

Four implementations refget currently exist across a diverse range of providers including ENA, Amazon Web Services (AWS) and Heroku (see supplementary materials). To test the compliance of these deployments, we developed a Python toolkit called the refget compliance documentation (https://compliancedoc.readthedocs.io/) and library suite (https://pypi.org/project/refget-compliance/). The compliance toolkit mandates each implementation will host three reference sequences; *Enterobacteria phage phiX174 sensu lato* (NC_001422.1) and *Saccharomyces cerevisiae S288C* chromosomes I (BK006935.2) and IV (BK006938.2). The compliance toolkit tests each server with a range of tests, with servers able to pass, fail or skip a test. This final option is used when a server is known to be unable to pass a test e.g., we do not test circular sequence retrieval if a server declares it does not support circular sequences. Tests are run daily against all known implementations and a HTML report is published at https://w3id.org/ga4gh/refget/compliance. We have also implemented a local Python interface in the *refget* Python package, hosted at PyPI (https://pypi.org/project/refget/). This package provides a local implementation of the refget protocol that can backed either by memory, SQLite, or MongoDB back-ends, and can connect to a remote API to provide local caching of retrieved results to improve performance for applications that require repeated lookups. It also provides Python functions to compute refget identifiers from within Python using raw sequences or FASTA files.

## DISCUSSIONS

Reference sequences remain a fundamental building block of bioinformatics providing a stable method of describing genomic events and annotation. Demand for reference sequences will only continue to increase as a result of the rise of genomic sequencing and analysis across multiple species, protocols and analysis methods. Refget formalises a method for generating identifiers from reference sequence and specifies an API to retrieve sequences, sub-sequences and metadata from said identifier. The specification is easy to implement with a mechanism to assert specification compliance. Refget can host any type of reference sequence, allows deployments to implement subsets of functionalities and provides a mechanism for deployments to programmatically declare this. We have also demonstrated how to deploy this specification across a variety of infrastructure. Our future plans will enable the definition of a reference sequence collection using checksums and sequence metadata, *e*.*g*., a genome, transcriptome or proteome and to provide a way to convert between known reference sequence names to digest based identifiers.

## ACNKOWLEDGEMENTS

The refget specification was developed as part of the Large Scale Genomics and Genomic Knowledge Standards work streams of GA4GH. We acknowledge the support of the GA4GH secretariat, data security work stream, regulatory and ethics work stream, steering committee and executive committee. We also acknowledge the advice and comments provided by Dixie Baker, James Bonfield, Gustavo Glusman, Reece Hart, John Marshall, Mike Love, Angel Pizzaro and Heidi Sofia.

