## supplementary material for "Refget: standardised access to reference sequences"

### Refget: standardised access to reference sequences - supplementary data

#### Supplementary material 1: Analysis of MGnify non-redundant protein database with TRUNC512 checksums

To ensure the suitability of our truncated SHA-512 based checksum representation, a collision analysis was performed on the May 2019 release of EMBL-EBI's MGnify peptide sequence database. Each FASTA formatted file was converted into a compressed CSV file detailing the sequence identifier, its TRUNC512 and MD5 checksums and the digested sequence. The resulting CSV files are reformatted to prioritise a checksum of interest, using awk, and then sorted lexicographically, using GNU sort. Sorting all sequence records in this way brings identical checksums together into neighbouring lines. The final file is then processed using a custom comparison script, which compares the current checksum record to the prior record. A collision is reported when two neighbouring lines in the final CSV file have the same checksum record. No TRUNC512 or MD5 collisions were detected in the 1,106,951,200 peptide sequences contained in MGnify's May 2019 data release.

Our analysis toolkit is available from https://github.com/andrewyatz/refget-application-note and can be used to analyse any collection of FASTA formatted sequences for collisions.

### Supplementary material 2: Description of refget implementations

The following document details four unique implementations of the refget protocol. Each section should be reviewed alongside any available source code and technical documentation, which will be referred to within this text.

#### Implementation 1: Refget reference implementation

| | |
|---|---|
| **Code availability** | Open source |
| **License** | Apache 2.0 |
| **Public repository** | https://github.com/andrewyatz/refget-server-perl |
| **Language** | Perl |
| **Deployment environment** | Traditional web and database environment. Pluggable sequence storage backend |
| **Support custom sequence loading** | Yes |

| | |
|---|---|
| **Supported checksum identifiers** | GA4GH identifier, MD5, TRUNC512 |
| **Public URL** | https://w3id.org/ga4gh/refget/reference |

Refget's reference implementation is implemented in Perl and is updated with each approved change to the refget specification. The server stores metadata (e.g. sequence length, known aliases) within a RDBMS such as PostgreSQL, MySQL or SQLite through the DBIx::Class (https://metacpan.org/pod/DBIx::Class) object relational mapper, implements its application logic using the Mojolicious framework (https://metacpan.org/pod/Mojolicious) and features an extensive library of tests. We recommend using PostgreSQL as the database and a Hypnotoad server behind a Nginx proxy should you wish to use the reference implementation in production.

Sequence storage is pluggable and can be delegated onto one of three solutions. First is a file system-based solution where sequences are stored by their TRUNC512 checksum. To avoid issues with file system limits files held in a directory, a hierarchy is created by taking hex character pairs from the TRUNC512 checksum and creating sub-directories. For example, a sequence with the TRUNC512 identifier `6681ac2f62509cfc220d78751b8dc524` would be held under the path `/66/81/6681ac2f62509cfc220d78751b8dc524`. This is a similar scheme employed by htslib to cache sequences on file systems, except htslib uses MD5 as the identifier. Databases or a redis store can also be used to hold sequences by a checksum identifier. The checksum employed by all indexing schemes can be configured using the application configuration file. The reference implementation is designed to be extensible and support additional alternative sequence storage solutions as required.

In addition, the application handles the GA4GH identifier format and can transparently convert requests for these into their equivalent TRUNC512 representations. This ensures backwards compatibility with the original SHA-512 based checksum algorithm.

#### Implementation 2: CRAM Reference Registry

| | |
|---|---|
| **Code availability** | Closed source |
| **License** | N/A |
| **Public repository** | N/A |
| **Language** | Java |
| **Deployment environment** | Traditional web environment, Oracle storage layer, Squid proxies |
| **Support custom sequence loading** | Partly. Hosts all sequences submitted and held in INSDC, so custom sequences must be submitted to ENA, Genbank or DDBJ. |
| **Supported checksum identifiers** | MD5 |
| **Public URL** | https://www.ebi.ac.uk/ena/cram |

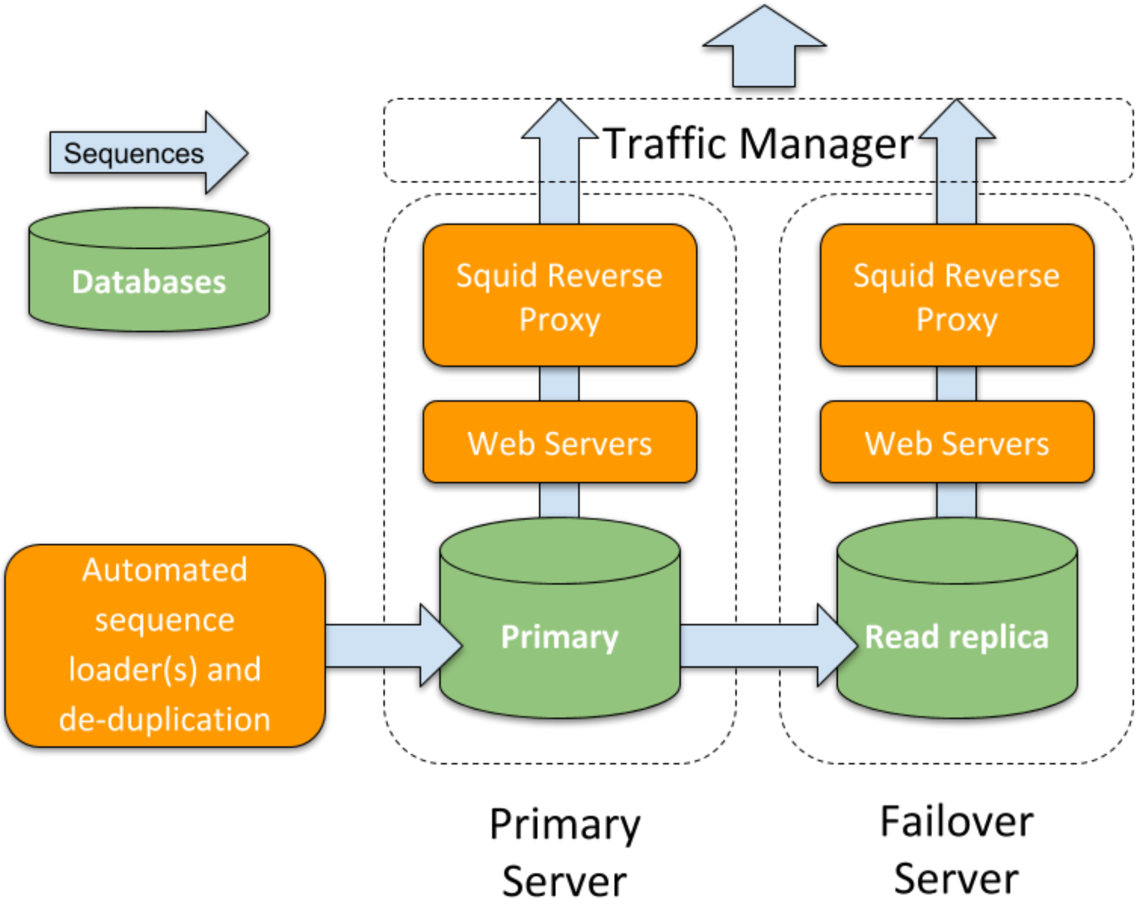


*Figure 1: ENA's CRAM Reference Registry architectural diagram. Sequences and metadata are stored in the backend Oracle database, which is replicated to a read-only instance. Two isolated stacks are made available via a traffic manager, each one fronted with a Squid proxy layer. This setup attempts to minimise service downtime and reduce load on the backend databases and web servers.*

ENA's CRAM Reference Registry is the original refget deployment serving CRAM's requirements for a centralised server for reference sequence retrieval. It served as the blueprint for the refget specification. All sequences held in the INSDC resources (ENA, Genbank, DDBJ) have been checksumed against the MD5 checksum, as noted in the CRAM specification. This server hosts over 2.5 billion sequences in an Oracle database. To reduce database and server load, the CRAM reference registry is hosted behind a Squid proxy server, to provide fast access to frequently accessed sequences.

#### Implementation 3: Container hosted on AWS

| | |
|---|---|
| **Code availability** | Open source |
| **License** | Apache 2.0 |
| **Public repository** | https://github.com/ga4gh/refget-cloud |
| **Language** | Python |
| **Deployment environment** | Refget protocol functions packaged as a Tornado web server, and deployed on Amazon Web Services (AWS). The containerized application is deployed through Amazon's Elastic Container Service (ECS) using the Fargate launch type. The application connects to a Public Dataset S3 bucket via HTTPS. |
| **Support custom sequence loading** | No |
| **Supported checksum identifiers** | GA4GH identifier, MD5, TRUNC512 |
| **Public URL** | https://w3id.org/ga4gh/refget/aws-container |

The Refget Cloud codebase serves as an extensible, generic library of refget functions that can be extended to work in a variety of cloud-based deployment contexts and using different cloud storage solutions. In this case, the refget service has been deployed as a containerized web server on AWS. Elastic Container Service (ECS) pulls the application's public docker image and launches containers via Fargate, which automatically provisions VM instances. The application sits behind an API Gateway, which enables SSL and passes the client request to the container. The container points to a public S3 bucket serving reference sequences from the INSDC consortium; the service passes incoming requests to S3, either redirecting the client directly to the S3 object or modulating the reference sequence before responding to the client. The deployment process uses AWS CloudFormation templates to achieve continuous delivery and infrastructure as code. A sample template is available in the repository. Figure 2 illustrates refget service architecture using AWS resources.

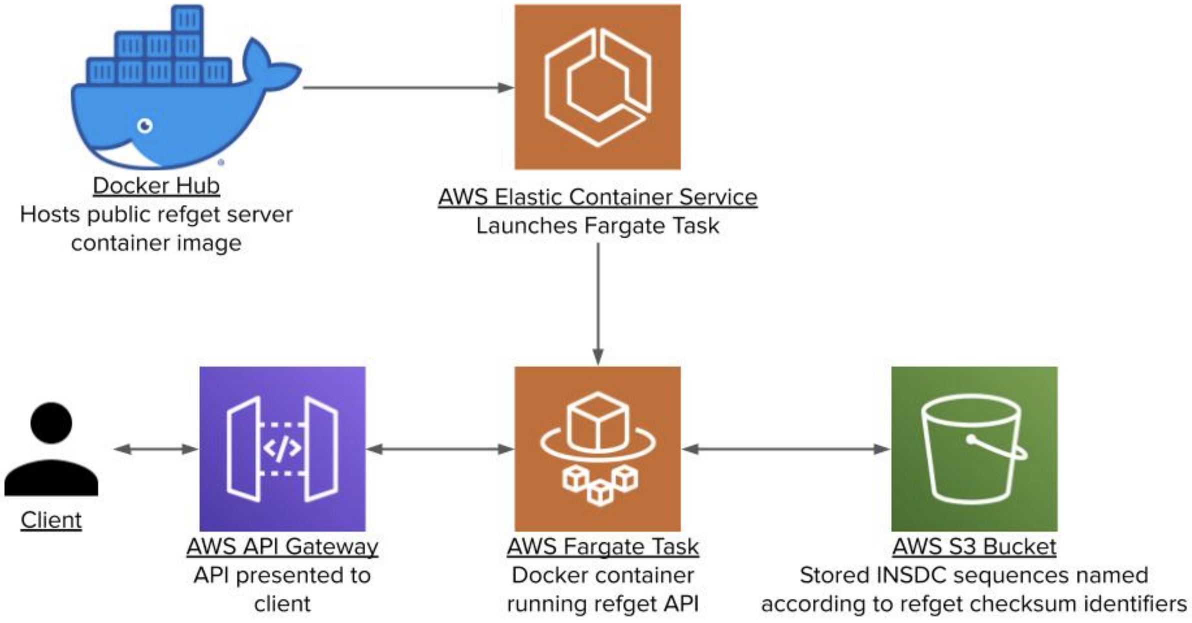


*Figure 2: Containerized refget service architecture using AWS cloud resources*

S3 provides an ideal storage solution for the refget service, as its properties allow it to handle most of the refget request directly. Overall, the refget container initially handles request parameter and header validation, and, if no errors are detected, offloads the responsibility of returning sequences and metadata to S3 via redirection. S3 can handle all metadata requests, full sequence requests, and subsequence requests specified via ‘Range’ header. Only when sub-sequences are requested via ‘start’ and/or ‘end’ query string parameters does the web server need to process the reference sequence directly.

S3 object properties allow sequences and metadata to be accessed by GA4GH, MD5, and TRUNC512 checksum identifiers without backend duplication of data, and without running a database. Sequences and metadata are stored in files named according to their GA4GH (primary) identifier. Empty files named according to a sequence’s MD5 and TRUNC512 (secondary) identifiers are also uploaded and affixed with a ‘Website-Redirection-Location’ metadata property that redirects file requests to the GA4GH identifier. Figure 3 illustrates this principle.

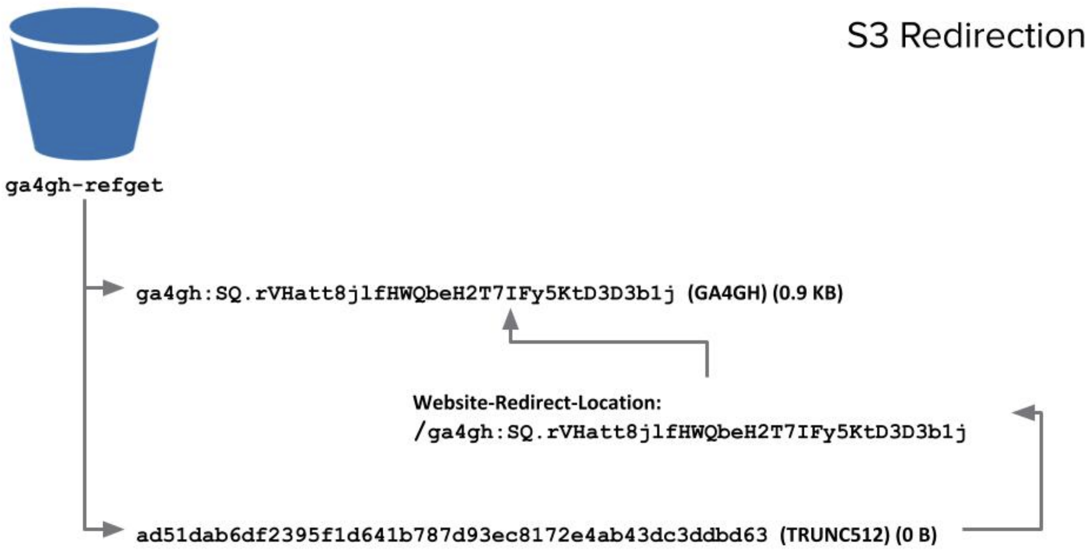


*Figure 3: Example redirection within S3. Incoming requests using the GA4GH identifier return the reference sequence stored under the corresponding file. Incoming requests using the TRUNC512 identifier are redirected to the GA4GH file, which ultimately returns the sequence. MD5 identifier requests also redirect to the GA4GH file.*

#### Implementation 4: Serverless refget on AWS

| | |
|---|---|
| **Code availability** | Open source |
| **License** | Apache 2.0 |
| **Public repository** | https://github.com/ga4gh/refget-cloud |
| **Language** | Python |
| **Deployment environment** | Refget protocol functions packaged as serverless, AWS lambda-compatible functions. The lambda functions sit behind an API Gateway and connect to a Public Dataset S3 bucket. |
| **Support custom sequence loading** | No |
| **Supported checksum identifiers** | GA4GH identifier, MD5, TRUNC512 |
| **Public URL** | https://w3id.org/ga4gh/refget/aws-serverless |

The same Refget Cloud codebase used in the above AWS ECS Fargate deployment has been repurposed to work in a serverless context. In this case, a refget service has been created without any provisioning of servers, VMs, or container instances. Instead, the service consists of three AWS Lambda functions (one for each endpoint in the refget specification); each function is associated with an HTTPS request route via API Gateway.

The serverless refget points to the same INSDC public S3 bucket. This service is best considered as a technical demonstration to complement the above containerized service; together they illustrate how refget can be deployed in multiple contexts using cloud resources. An AWS CloudFormation template for deploying serverless refget is also available in the GitHub repository. Figure 4 illustrates serverless refget's simple architectural design.

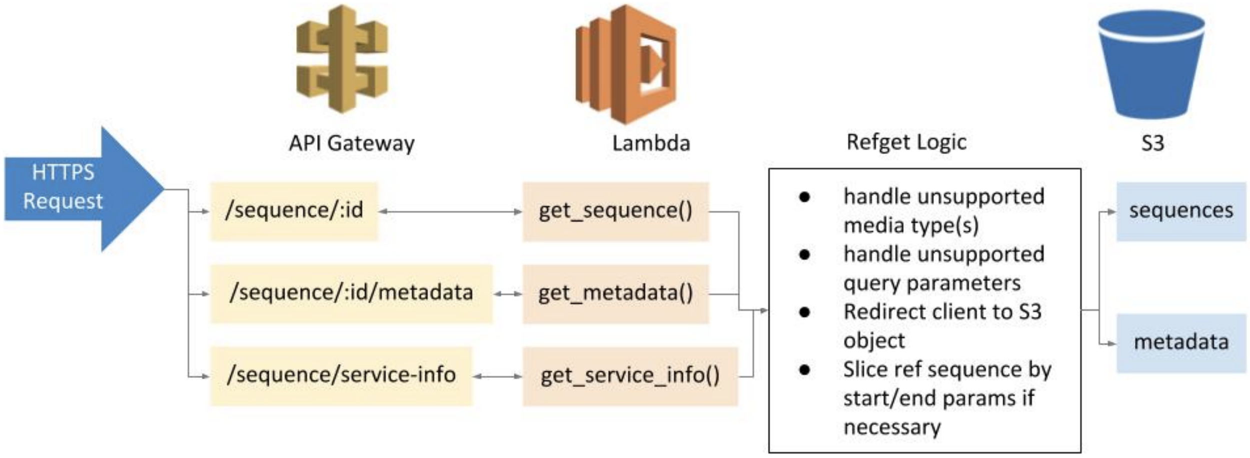


*Figure 4: Serverless refget deployment. API Gateway maps HTTPS request routes to lambda functions in the python library. The functions perform initial refget validation logic before offloading returning sequences and metadata to the S3 bucket via redirection.*

### Supplementary material 3: Clients

#### Local Refget Python Package

| | |
|---|---|
| **Code availability** | Open source |
| **License** | BSD 2 |

| | |
|---|---|
| **Public repository** | https://github.com/refgenie/refget |
| **Language** | Python |
| **Deployment environment** | A local Python package that provides a Python API client that can be backed either by local storage, or by a remote refget API. |
| **Support custom sequence loading** | Yes |
| **Supported checksum identifiers** | GA4GH identifier, MD5, TRUNC512 |
| **Public URL** | N/A (Local use only) |

The refget Python package is not intended to create an HTTP(s)-service like the above implementations, but to provide a user-side Python client to simplify interacting with such services. Users may use the Python package to easily query existing remote refget implementations and retrieve sequences from within Python. The package provides the ability to store retrieved sequences in a local database, and also to add new sequences to such a database, which may be backed by an instance of SQLite, MongoDB, or simple Python dict objects to store sequences in memory. This effectively provides local caches of sequences which can improve lookup performance. The package also enables users to easily set up a custom, local sequence store for use within Python that can mix public and private sequences. Furthermore, the Python package may be used independently of any remote instance if sequences are all provided locally. It also provides other related helper functions that simplify working with refget servers or creating sequence identifiers.
